# Global characterization of gene expression in the brain of starved immature *R. prolixus*

**DOI:** 10.1101/2022.09.01.506236

**Authors:** Jessica Coraiola Nevoa, Jose Manuel Latorre-Estivalis, Fabiano Sviatopolk-Mirsky Pais, Newmar Pinto Marliére, Gabriel da Rocha Fernandes, Marcelo Gustavo Lorenzo, Alessandra Aparecida Guarneri

## Abstract

Background: *Rhodnius prolixus* is a vector of Chagas disease and has become a model organism to study physiology, behavior, and pathogen interaction. The publication of its genome allowed initiating a process of comparative characterization of the gene expression profiles of diverse organs exposed to varying conditions. Brain processes control the expression of behavior and, as such, mediate immediate adjustment to a changing environment, allowing organisms to maximize their chances to survive and reproduce. The expression of fundamental behavioral processes like feeding requires a fine control in triatomines because they obtain their blood meals from potential predators. Therefore, the characterization of gene expression profiles of key components modulating behavior in brain processes, like those of neuropeptide precursors and their receptors, seems fundamental. Here we study global gene expression profiles in the brain of starved *R. prolixus* fifth instar nymphs by means of RNASeq sequencing. Results: The expression of neuromodulatory genes such as those of precursors of neuropeptides, neurohormones, and their receptors; as well as the enzymes involved in the biosynthesis and processing of neuropeptides and biogenic amines were fully characterized. Other important gene targets such as neurotransmitter receptors, nuclear receptors, clock genes, sensory receptors, and takeouts were identified and their gene expression analyzed. Conclusion: We propose that the set of neuromodulation-related genes highly expressed in the brain of starved *R. prolixus* nymphs deserves functional characterization to allow the subsequent development of tools targeting them for bug control. As the brain is a complex structure that presents functionally-specialized areas, future studies should focus on characterizing gene expression profiles in target areas, e.g. mushroom bodies, to complement our current knowledge.

## Introduction

Triatomines are hematophagous insects that can transmit *Trypanosoma cruzi*, the etiological agent of Chagas disease. It is estimated that this neglected disease affects 7 million people, located mostly in Central and South America. The study of their biology is relevant because *T. cruzi* transmission is mostly controlled by eliminating domiciliated bugs (1).

Triatomines are nocturnal insects that assume an akinetic state while hidden in shelters during daylight hours. At nightfall, they eventually start non-oriented locomotor activity, outside shelters to search for hosts. For host recognition, starved bugs detect cues released by vertebrates, such as radiant heat, water vapor, carbon dioxide, and other odorants (2). The decision to leave a shelter and engage in foraging is risky, as triatomine hosts are often predators as well. For this reason, starved bugs mostly leave the protection of the shelters when a robust set of host clues is present (3–5). Bugs of all nymphal instars and adults of both sexes feed on blood and can tolerate long starvation. Whereas nymphs have to feed to be able to molt, adult females require nutrients to produce eggs (6). Starved insects orientate towards host-emitted stimuli, while fed insects can remain indifferent or avoid these cues depending on the time elapsed after feeding (7).

The central nervous system (CNS) is the main regulator of physiology and behavior. Besides processing sensory information, the brain is the major accumulation of neuropiles integrating neural activity of sensory, memory, and proprioceptive nature (8). As such, it has a main role in the coordination of motor responses, adjusting their proper timing through a set of clock neurons (9–11). Signal transfer and modulation of neural processes in the CNS depend on neuroactive compounds, including neurotransmitters of diverse chemical nature like biogenic amines and neuropeptides, and their receptors (12,13). Neuropeptides and biogenic amines can also act as endocrine factors mediating signaling processes in multicellular organisms, and in the case of insects, they are fundamental in coordinating growth and development, as well as physiological processes such as metabolism, diuresis, digestion, reproduction, and behavior (14,15). *R. prolixus* has been widely used as a model for insect physiology studies, including research on reproduction, development, immunology, and vector-parasite interactions (16). After the publication of the *R. prolixus* genome sequence (17), and the introduction of next-generation sequencing (NGS) methods, several studies have described genetic and molecular components underlying the physiology of *R. prolixus* (14,15,18–26). Transcriptomic studies allowed the discovery of new genes and transcripts, the identification of differentially expressed genes, determining targets for broader functional analyses. Some transcriptomes have analyzed gene expression in different *R. prolixus* tissues such as salivary glands (27), ovaries (28,29), gut (25), testicles (30), and antennae (22). Furthermore, this technique allowed defining the molecular bases of female reproductive physiology under differing nutritional states (18), as well as characterizing the innate immune system of these bugs at the molecular level (31). Several of these bioinformatic analyses have recently shown that the genome of *R. prolixus* has many missing or miss-annotated genes (22,29,31), highlighting the importance of transcriptomes for improving the quality of the annotated genome of *R. prolixus*. Therefore, the present study aims to describe the genetic components that serve as the molecular neural bases controlling behavior in the brain of unfed *R. prolixus* nymphs.

## Materials and methods

### Insects

*Rhodnius prolixus* were obtained from a colony derived from insects collected in Honduras around 1990 and maintained by the Vector Behavior and Pathogen Interaction Group at the René Rachou Institute, Belo Horizonte, Brazil. Insects were monthly fed on citrated rabbit blood obtained from CECAL (Centro de Criação de Animais de Laboratório, FIOCRUZ, Rio de Janeiro, Brazil) offered through an artificial feeder at 37 °C, alternating with feeding on anesthetized chicken and mice. Chickens were anesthetized with intraperitoneal injections of a mixture of ketamine (20 mg/kg; Cristália, Brazil) and detomidine (0.3 mg/kg; Syntec, Brazil), and mice with ketamine (150 mg/kg; Cristália, Brazil) and xylazine (10 mg/kg; Bayer, Brazil). The procedures using animals were approved under license number LW-61/2012 (Committee for Ethics in the Use of Animals, CEUA-FIOCRUZ). Insects were reared in the insectary under 27 ± 2 °C, 51 ± 7% of relative humidity and natural illumination.

### RNA extraction and Illumina sequencing

Thirty-day-old 5^th^ instar nymphs were used in this experiment. Insect brains were dissected on a freeze cold dissecting dish (BioQuip, Gardena, CA, US), collected with forceps, and immediately transferred to a microtube immersed in dry ice and added 1 mL of TRIzol™ reagent (Invitrogen, Thermo Fisher Scientific, MA, USA). For sample completion, dissections occurred along three days, exclusively between 2 and 4 PM. Six independent replicates were performed, each composed of a pool of 20 brains. RNA extraction was performed with TRIzol™ according to the manufacturer’s instructions. Total RNA concentrations were determined using a Qubit 2.0 Fluorometer (Life Technologies, Carlsbad, CA, US). The libraries were constructed using the TruSeq Stranded mRNA Sample Preparation Kit (Illumina, San Diego, CA) and sequenced on an Illumina HiSeq 2500 platform at the Max Planck Genome Center in Cologne (Germany). Approximately 15 million reads were obtained for each library, using 150 base-pair paired-end reads. The raw sequence dataset is available with the NCBI-SRA Bioproject number PRJNA853796 at NCBI.

### Bioinformatic analysis

Raw reads were filtered and trimmed for low-quality bases using Trimmomatic (v0.36) (32), according to standard quality score parameters (Phred-33 (>15); and 50 base-pair minimum length). Then, STAR v2.6.0 (33) was used with default parameters to map reads to the *R. prolixus* reference genome (version RproC3.3) accessed through the VectorBase website (34). Mapped reads were assigned to each gene through BEDTools coverage (v2.29.2) based on an updated gene annotation file (22). Gene length of mapped reads and total counts were used for calculating Fragment *per* Kilobase *per* Million mapped reads (FPKM) values for target genes in each library. Subsequently, target gene expression (as Log10 (FPKM+1)) was depicted in heatmaps built using the pheatmap R package (v1.0.12). Finally, all genes were ranked according to the highest expression using Log 10 (FPKM+1) values. Finally, all genes were ranked according to the highest expression using Log10 (FPKM+1) values, and from the genes with the 100 highest scores, the top 50 that were present in at least four of the six libraries, and which presented annotation were selected. The identity and putative functions of these highly expressed genes were obtained from BLASTp searches against NCBI (including only insect sequences) and InterPro description from VectorBase (34). Genes with sequences shorter than 100 amino acids and/or without functional information were not included in this ranking.

## Results and discussion

### Overall analysis

RNA-Seq data from starved 5^th^ instar *R. prolixus* brain transcriptomes were summarized in Table 1. After filtering and trimming raw reads, all libraries showed coverage of at least 13 million reads. The number of uniquely mapped reads against the *R. prolixus* genome ranged from 8,8 M to 10,3 M reads.

**Table 1.** Summary of RNA-Seq metrics from *R. prolixus* brain transcriptomes.

| Sample name | Raw reads | Clean reads | Uniquely mapped | Uniquely mapped (%) |
| --- | --- | --- | --- | --- |
| Brain_rep1 | 15,816,907 | 12,843,740 | 9,235,291 | 71.90 |
| Brain_rep2 | 16,109,824 | 13,304,387 | 8,860,721 | 66.60 |
| Brain_rep3 | 16,379,827 | 13,237,314 | 9,795,612 | 74.00 |
| Brain_rep4 | 16,367,691 | 13,094,877 | 9,451,882 | 72.18 |
| Brain_rep5 | 16,811,723 | 13,819,788 | 9,699,418 | 70.18 |
| Brain_rep6 | 16,462,326 | 14,243,670 | 10,330,221 | 72.52 |

**Sample name:** name of replicate; **Raw Reads:** original sequencing reads; **Clean Reads:** number of reads after filtering; **Uniquely** **mapped:** number of reads that were uniquely mapped to the reference genome; **Uniquely mapped (%):** percentages of uniquely mapped reads.

Based on our brain library outputs, six lists of 100 top expressed genes were obtained after ranking their Log10 (FPKM+1) values. The comparison of these ranks allowed building a consensus list depicting genes that ranked top 50 in at least four out of six libraries (Table 2). Several initiation and elongation factors together with ribosomal proteins were identified among top expressed genes, probably reflecting protein biosynthesis induced by starvation. Several heat shock proteins presented high expression in the brain, also probably due to starvation-generated stress. Different types of soluble carrier proteins, like odorant binding proteins, takeouts and lipocalins were very abundantly expressed in the CNS, suggesting roles other than odor transportation or detection. Finally, other highly expressed genes were the neuroendocrine secretory protein 7B2 which functions as a specific chaperone for the prohormone convertase 2 (PC2) (35), an enzyme required for the maturation of neuropeptide and peptide hormone precursors; and the Glutamine synthetase that catalyzes the synthesis of glutamine, which has a central role in nitrogen metabolism and the regulation of neurotransmitter production (36).

**Table 2.**
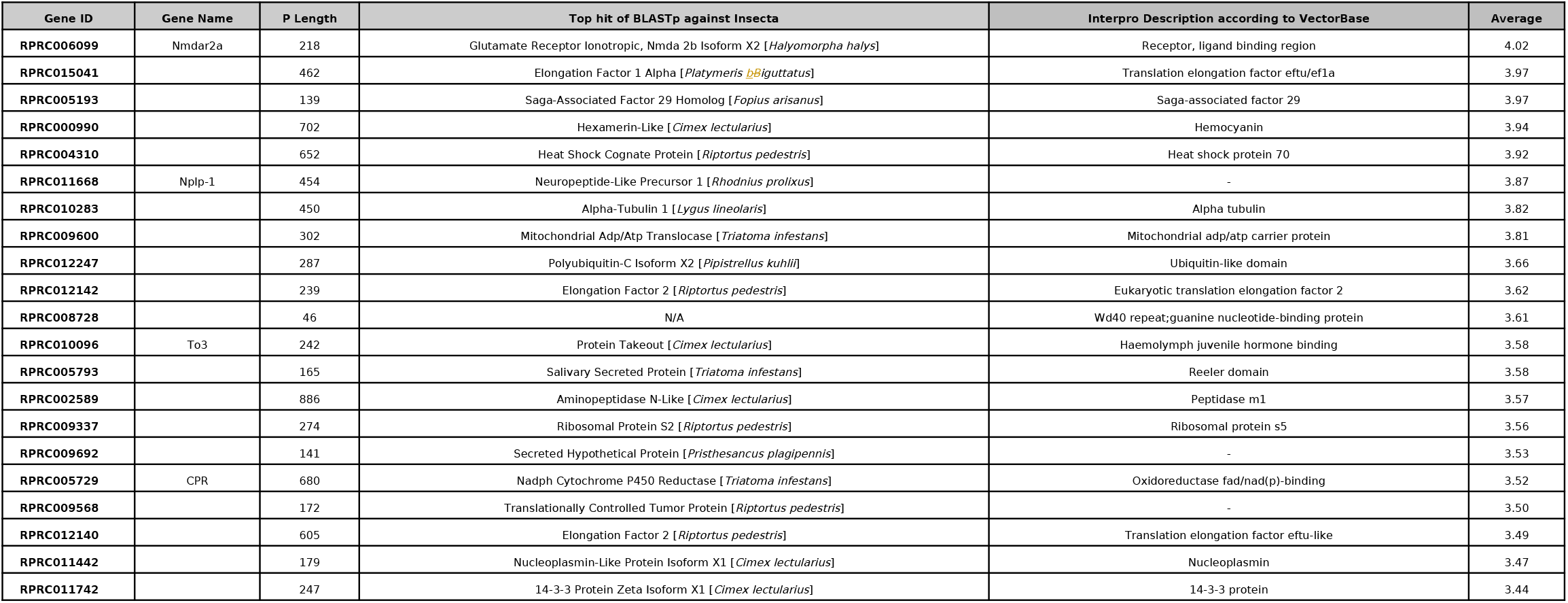
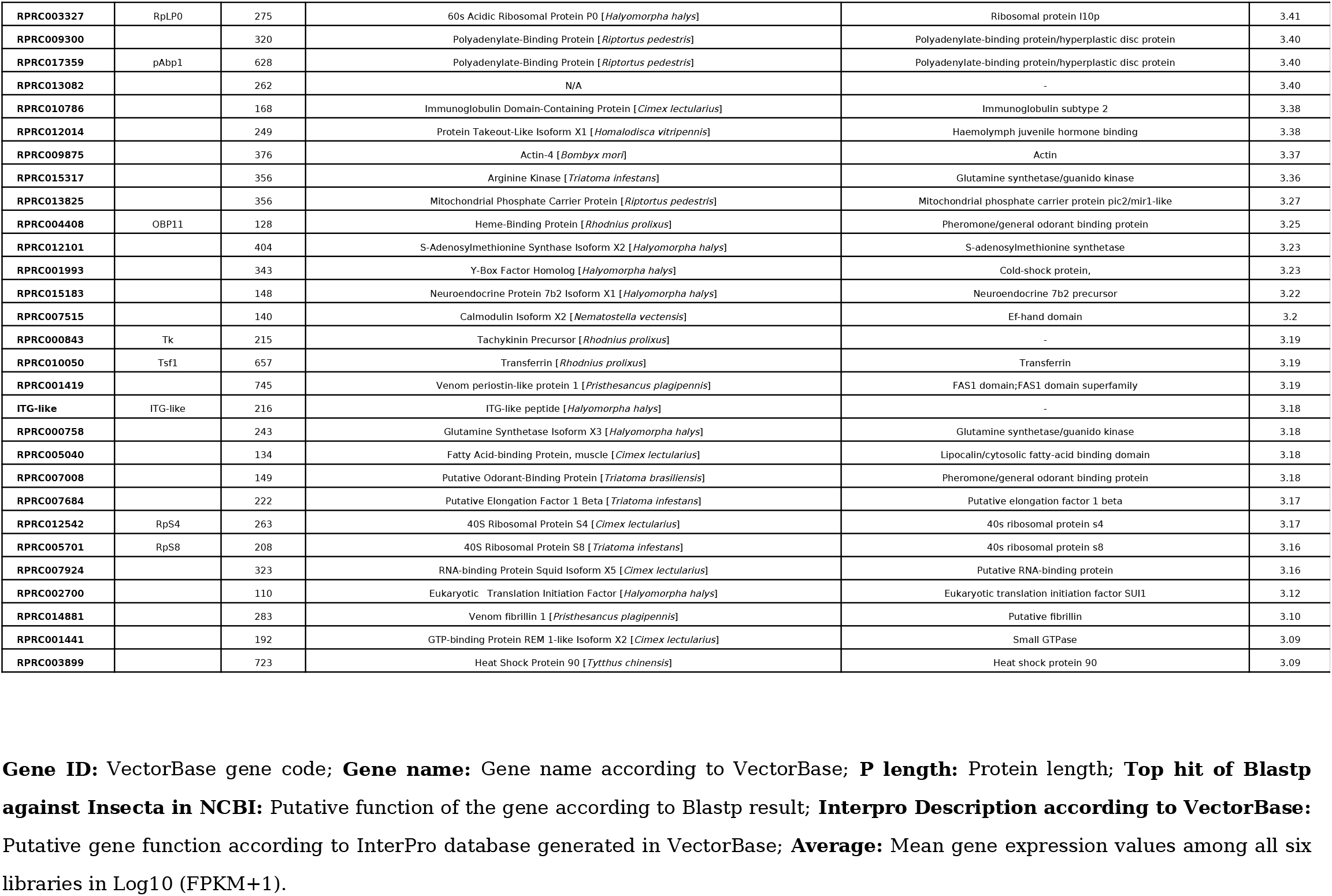
Consensus list of the top 50 most highly expressed genes in brain transcriptomes of starved *R. prolixus*.

### Neuropeptide precursor genes

Neuropeptide precursor gene (NPG) expression patterns in insects tend to be stereotyped, and each neuropeptide may be involved in neurotransmission, in synaptic neuromodulation or in conveying neuroendocrine signals at peripheral targets (12). Currently, structural and functional studies on insect neuropeptides are being performed to develop new insect control approaches. This is especially true for neuropeptides involved in developmental, nutritional, and survival processes (24). For this reason, we focussed on the neuropeptides showing highest expression in our study.

The four most highly expressed NPGs (Neuropeptide-like precursor 1 - NPLP1, Tachykinin - TK, ITG and NVP-like) presented expression values > 3 (Log10 (FPKM +1)). Most genes coding for neuropeptides (25 genes) had expression values between two and three, while nine had values between 1 and 2. As expected for some of them, ecdysis triggering hormone (ETH), elevenin 1 (Ele1), eclosion hormone (EH), sulphakinin (SK), and sifamide (SIFa) had low expression values < 1. Adipokinetic hormone/corazonin-related peptide (ACP) was the only NPG out of 44 annotated for R. prolixus (22) that was not expressed in any of the six libraries here analyzed (Fig 1a).

**Fig 1.**
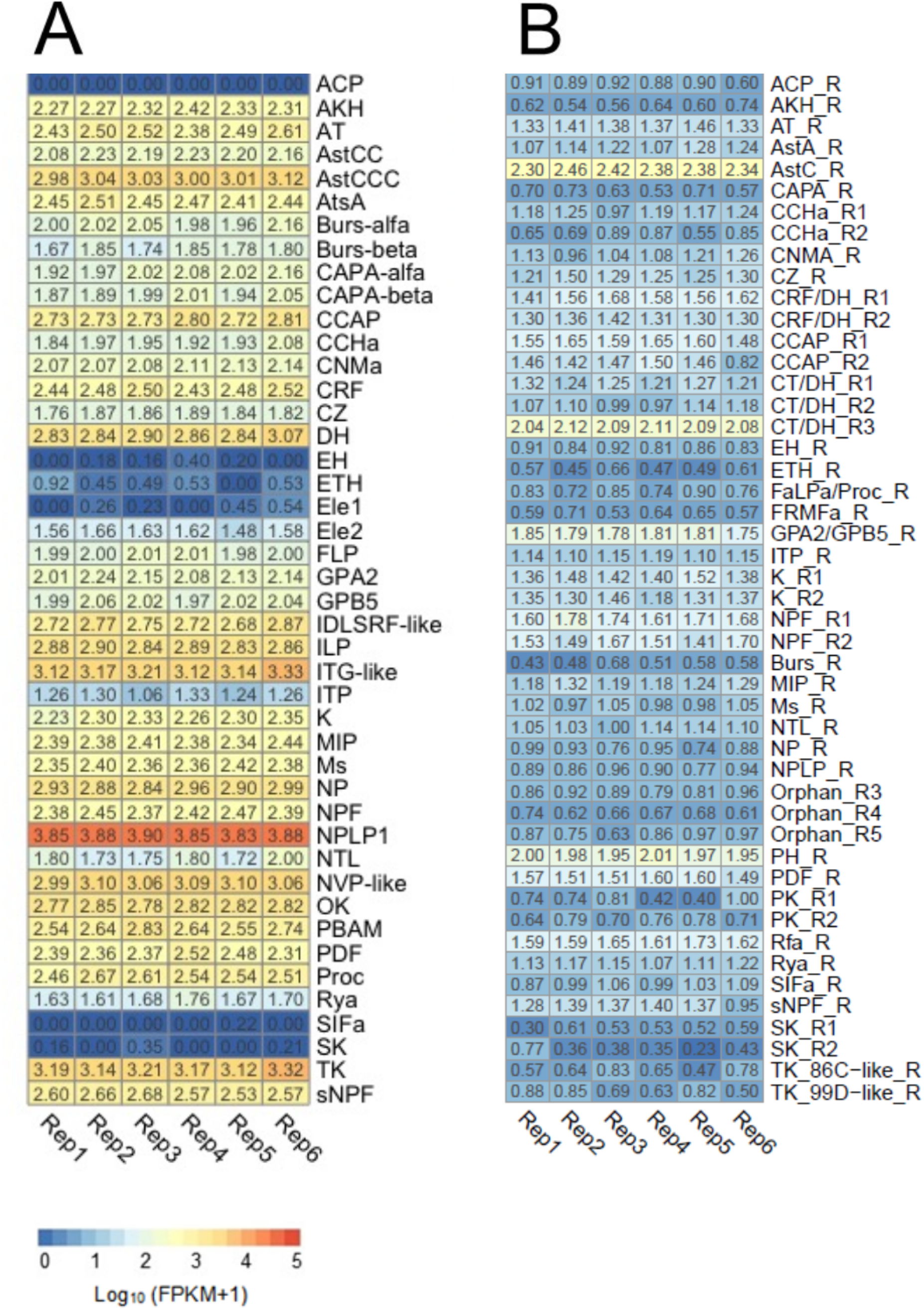
Expression profiles of neuropeptide precursor and neuropeptide receptor genes. (A) Heat map depicting the expression level of neuropeptide precursor genes, and (B) neuropeptide receptor genes in the brain of *R. prolixus* nymphs. Expression (displayed as Log10 (FPKM+1)) is represented by means of a color scale in which blue/orange represent lowest/highest expression. Each column represents the expression of one library.

Neuropeptide-like precursor 1 (NPLP1) was the NPG showing the highest expression in our transcriptomic database. Furthermore, it ranked amongst the 10 most highly expressed genes in the brain, presenting a mean of 3.86 ± 0.01 Log10 (FPKM+1) among our libraries. There were two different isoforms with distinct expression patterns detected previously in *R. prolixus*, isoform A expressed in CNS, ovaries, testes, and antennae, while expression of isoform B was only detected in antennae. The presence of mature peptides from NPLP1 precursors was also previously detected in the salivary glands of *R. prolixus* using proteomics (26). The high expression observed for this neuropeptide precursor in our work suggests a relevant role in the CNS. Indeed, Sterkel et al. (26) showed that the levels of two neuropeptides encoded by the NPLP1 precursor gene significantly decreased in the CNS 24 hours after feeding, suggesting a role connected to starvation. A decrease in NPLP1 expression was also observed in the antennal lobe of mosquitoes after blood or sugar ingestion (37).

According to our brain dataset, tachykinin (TK) was the NPG showing the second-highest expression level (3.19 ± 0.02 Log10 FPKM +1). Sterkel et al. (26) have also detected TK in the CNS of *R. prolixus* by means of peptidomics, not observing changes in its abundance after blood ingestion. This NPG presents expression in a wide set of *R. prolixus* tissues, as its transcripts have been th th detected in salivary glands, fat body, dorsal vessel, intestinal tract of 5^th^ instar nymphs (by RT-qPCR) (38) and in the antennae of 5^th^ instar nymphs and adults (using RNA-Seq) (22). Studies on other insects have shown that TKs can modulate early olfactory processing at the olfactory lobes, circuits controlling locomotion and food search, aggression, metabolic stress and nociception (39). Similar roles could be expected in triatomines based on the high expression of the TK gene observed in our study.

ITG-like and NVP-like genes also presented high expression values (3.18 ± 0.03 and 3.06 ± 0.01 Log10 (FPKM +1), respectively) in our database; nonetheless, to our knowledge, a functional characterization is not available for these peptides so far. Latorre-Estivalis et al. (22) detected high expression of ITG-like in the antennae of immature and adult *R. prolixus* and proposed a modulatory role for this neuropeptide at the peripheral level. In its turn, Leyria et al. (18) indicated that blood ingestion induced ovarian down-regulation of the NVP-like gene, suggesting a role in *R. prolixus* reproduction (18). The quantitative peptidomic analysis of the *R. prolixus* CNS by Sterkel et al. (26) showed that the abundance of both neuropeptides significantly decreases a few hours post-blood meal, indicating their implication in a neuroendocrine response to feeding. Considering this and the very high expression observed for ITG-like and NVP-like in our dataset from starved bugs, we suggest that they might act by signaling starvation status. Functional genetic studies based on gene silencing should be implemented to verify potential behavioral phenotypes that could offer evidence about the roles of these neuropeptides in *R. prolixus*.

Insulin-like peptide (ILP) presented a high expression in our database (2.86±0.01 Log10 FPKM +1). This result was consistent with the characteristics of our samples (brain from starved bugs) and the putative function of this neuropeptide as a modulator of lipid and carbohydrate metabolism (40,41). Indeed, *in vitro* immunofluorescence studies in *R. prolixus* brains demonstrated strong and abundant fluorescence of ILP neurons in unfed nymphs, followed by an acute decrease 4 hours after ingestion of a food meal, indicating transport and release of these signaling molecules into the hemolymph soon after feeding (42). The high expression of this neuropeptide in the CNS of unfed insects was also observed by Leyria et al. (18). Interestingly, these authors observed that blood ingestion did not affect ILP gene expression in the CNS of *R. prolixus* (18).

### Neuropeptide processing enzymes

Mature neuropeptides are synthesized by a series of enzymatic steps that sequentially cleave and modify larger precursor molecules. One step of neuropeptide biosynthesis involves peptide amidation, a process that occurs on half of the known bioactive neuropeptides (43,44). The expression pattern observed for these enzymes might be a good proxy to estimate their abundance and activity (45), deserving their subsequent functional characterization.

A set of eleven neuropeptide processing enzymes was previously annotated for *R. prolixus* (22), using sequences of *D. melanogaster* as references (46). All of these enzymes showed relative expression values higher than 1 (Log10 (FPKM+1)) (S1 Fig). Peptidyl alpha hydroxyglycine alpha amidatinglyase2 (PAL2) and peptidylglycine α-amidating monooxygenase (PHM), the enzymes involved in amidation reactions, were the most highly expressed genes of this set (PAL2 – 2.89±0.02, PAM – 2.74±0.05). The brain expression pattern shown by this group of genes seems to differ when compared to that shown in an antennal transcriptome of *R. prolixus* (22).

### Neuropeptide receptors

Whether a tissue is targeted by a certain neuropeptide is defined by the presence of the corresponding neuropeptide receptor on the surface of its cells. Neuropeptide receptor genes showed much lower expression values than neuropeptide gene precursors. Only two out of the 48 neuropeptide receptor genes analyzed showed a relative expression higher than 2 (Fig 1b); AstC receptor with 2.38±0.02 Log10 (FPKM +1) and calcitonin-like diuretic hormone receptor 3 -CT/DH-R3 with 2.08±0.01 Log10 (FPKM +1). Twenty-six genes (54%) presented a relative expression between 2 and 1, and twenty-two genes (45%) values lower than 1.

The expression of the AstC receptor in the *R. prolixus* CNS had been previously reported by Ons et al. (15) using RNA-Seq. These authors did not see expression changes for this receptor at the CNS after blood ingestion. Furthermore, Villalobos-Sambucaro et al. (47) observed the presence of this receptor in the hindgut, midgut and dorsal vessel, and showed that the receptor and its ligand play a key myoregulatory and cardioregulatory role in *R. prolixus*. Our results suggest that the AstC receptor may also have a fundamental role at the central level. Two receptors for CT/DH (named R1 and R2) were previously described in *R. prolixus*, their transcript expression being detected in the CNS and reproductive tissues (48). The existence of a third CT/DH receptor in *R. prolixus* was suggested by Ons et al. (15) and confirmed by an antennal transcriptome (22). The latter showed that this receptor had increased expression in male antennae when compared to those of nymphs, suggesting a sexually dimorphic role for this peptide (22). Our results suggest that this receptor may also act at the central level.

### Neurotransmitter receptors

As expected, neurotransmitter receptors tended to present higher expression values than the other receptor gene families studied here. N-methyl-D-aspartate receptor type 2A (NMDAr2a) was the gene with the highest expression in the whole brain transcriptome (Fig 2; 4.02±0.01 Log10 (FPKM +1)). Muscarinic acetylcholine receptor type A (AChR-A) and dopamine 1-like receptor 1 (DOP1) were the other two genes that presented FPKM values higher than 2 in this gene set (2.12±0.05 Log10 (FPKM +1) and 2.11±0.007 Log10 (FPKM +1) for AChR-A and DOP1, respectively). Only 5 hydroxytryptamine (serotonin) receptors 1 (5HT-1A-R) and 7a (5HT-7A-R) showed values lower than 1 (0.59±0.03 and 0.79+0.03 Log10 (FPKM +1) for 5HT-1A-R and 5HT-7A-R, respectively).

**Fig 2.**
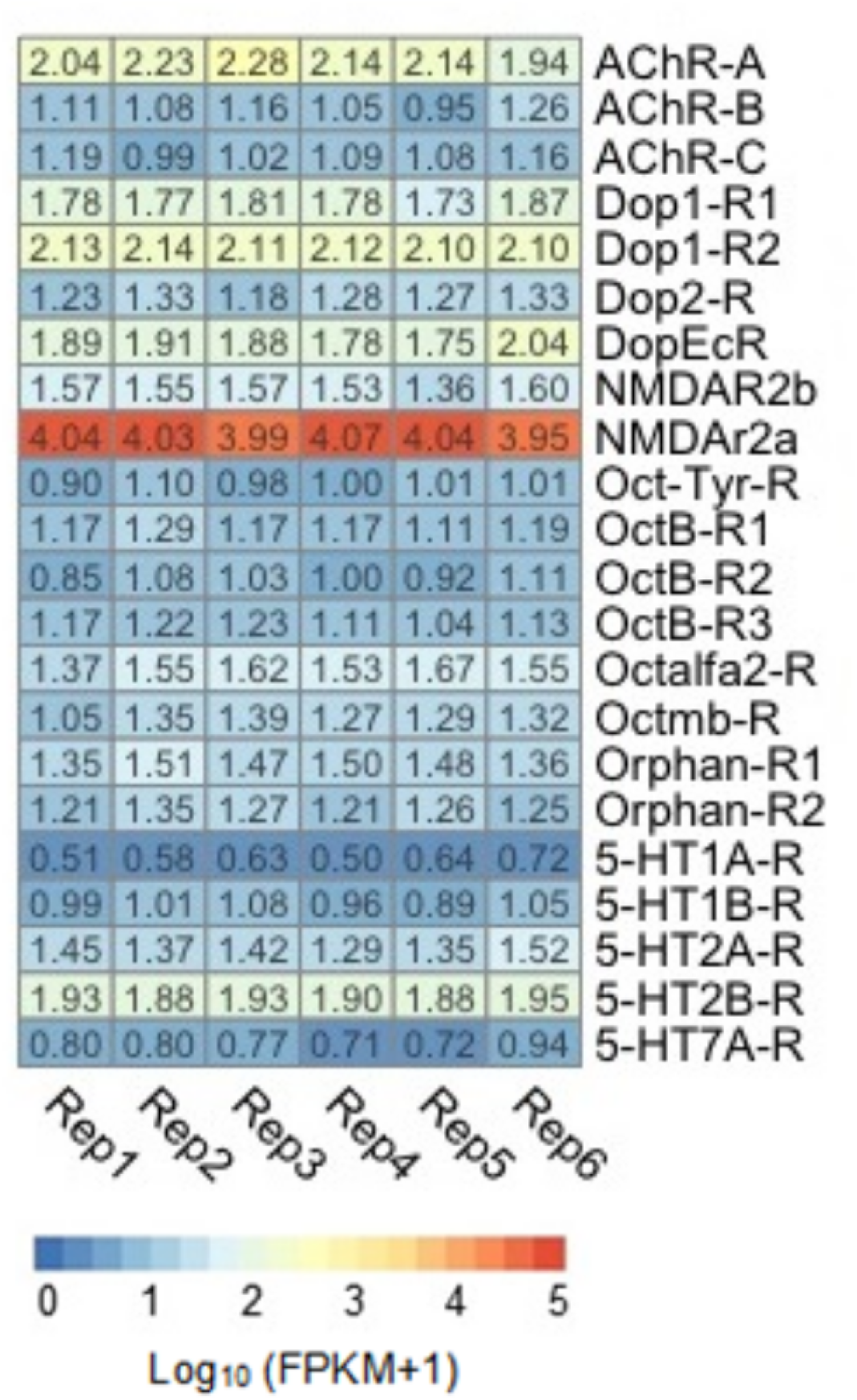
Expression profiles of neurotransmitter receptors genes. Heat map depicting the expression level of neurotransmitter receptor genes in the brain of *R. prolixus* nymphs. Expression level (displayed as Log10 (FPKM+1)) is represented by means of a color scale in which blue/orange represent lowest/highest expression. Each column represents the expression of one library.

N-methyl-D-aspartate (NMDA) receptors are one of the subtypes of ionotropic receptors that bind to L-glutamate, mediating an excitatory activity in the CNS of insects. The NMDARs are usually constituted of two subunits NR1 and NR2 (49). Even though these receptors were characterized in the brain of several invertebrate species, their functions in insects are poorly understood. However, their involvement in behavioral plasticity is already known (50,51). Similarly, a study of the evaluation of NMDAR expression in different tissues from female *Dactyola punctata*, showed that DpNR1A, DpNR1B and DpNR2 were highly expressed in the brain (52). In *Drosophila melanogaster*, both NMDA receptors called *dNR1* and *dNR2* were weakly expressed throughout the entire brain, with higher expression observed in some scattered cell bodies (53). As far as we know, this is the first report on the expression of this receptor in the brain of *R. prolixus;* its high expression suggesting a very relevant role in the neural physiology of these insects. Immunostaining experiments to characterize brain neuropiles depicting NMDAr2a expression will be required to initiate functional studies to uncover its putative function.

### Nuclear receptors

Most nuclear receptor genes presented low expression in the brain of unfed nymphs (Fig 3). Out of this gene set, the ecdysone induced protein 75B (*Eip75B*) receptor was the only gene that presented FPKM values higher than 2 (2.2±0.06 Log10 (FPKM +1). Nine out of 23 genes (39%) presented values between 1 and 2 Log10 (FPKM +1). The Eip75B and HR51 transcripts (the latter also known as *unfulfilled*) have been identified in central clock cells of *D. melanogaster* and control the expression of clock genes, playing an important role in the maintenance of locomotor rhythms (54–56). Similar roles could be proposed for *R. prolixus*, however, functional information is not available for *Eip75B* and *HR51* in this species; only Latorre-Estivalis et al. (22) reported similar expression values for the *Eip75B* gene in kissing bug antennae.

**Fig 3.**
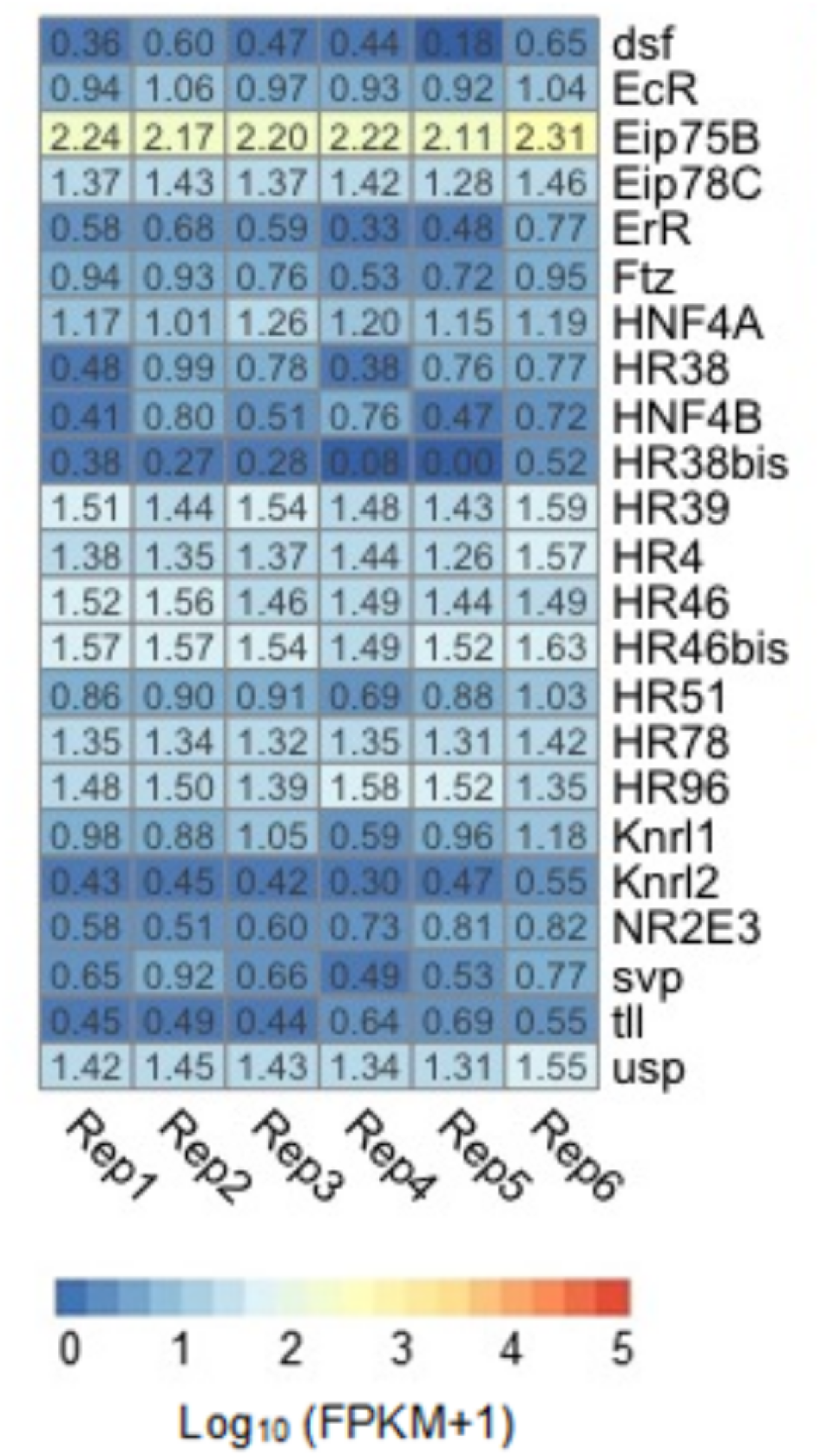
Expression profiles of nuclear receptors genes. Heat map depicting the expression level of nuclear receptor genes in the brain of *R. prolixus* nymphs. Expression level (displayed as Log10 (FPKM+1)) is represented by means of a color scale in which blue/orange represent lowest/highest expression. Each column represents the expression of one library.

### Clock and behavior-related genes

Clock genes are responsible for controlling circadian rhythms, and they can cycle in a synchronized way according to daily oscillations of environmental cues such as light and temperature (57). Even though most available information on clock gene function has been generated using *D. melanogaster* as a model, their fundamental roles and their high level of sequence homology suggest that their functions should be conserved across insect orders. A total of 31 clock genes have been previously described and annotated in the *R. prolixus* genome (17); however, as far as we know, this is the first time that the expression of the whole clock gene set is studied in this insect. As all our brain samples were generated at the same interval of the daily cycle (2-4 PM), the expression profiles obtained define the levels of expression of clock genes at this time. Except for *vrille* (*vri*), *cycle* (*cyc*), *single-minded* (*sim*) and *timeless* (*tim*) genes, the rest of the clock genes had expression higher than 1 Log10 (FPKM+1) values (Fig 4). The most highly expressed genes (> 2 Log10 (FPKM+1)) were: casein kinase 2 (*ck2*), protein phosphatases (*Pp*) 1a and 2a, *no circadian temperature entrainment* (*nocte*), and poly(A) binding proteins (*pAbp*) 1 and 2. These genes showing high expression probably have key roles in the brain of unfed kissing bugs. Based on *D. melanogaster* studies, *pAbps* genes form a complex with *twenty-four* (*tyf*) and *Ataxin-2* (*Atx2*) that maintain circadian rhythms in locomotor behavior (58). The *Pp2A* gene, which is also highly expressed, controls the cyclic expression of the PER protein (59), which was also detected in our dataset. The *nocte* gene encodes a protein involved in temperature compensation of the circadian clock in *Drosophila*. It would be relevant to characterize 24h expression profiles of clock genes in the *R. prolixus* CNS to identify genes with cycling expression.

**Fig 4.**
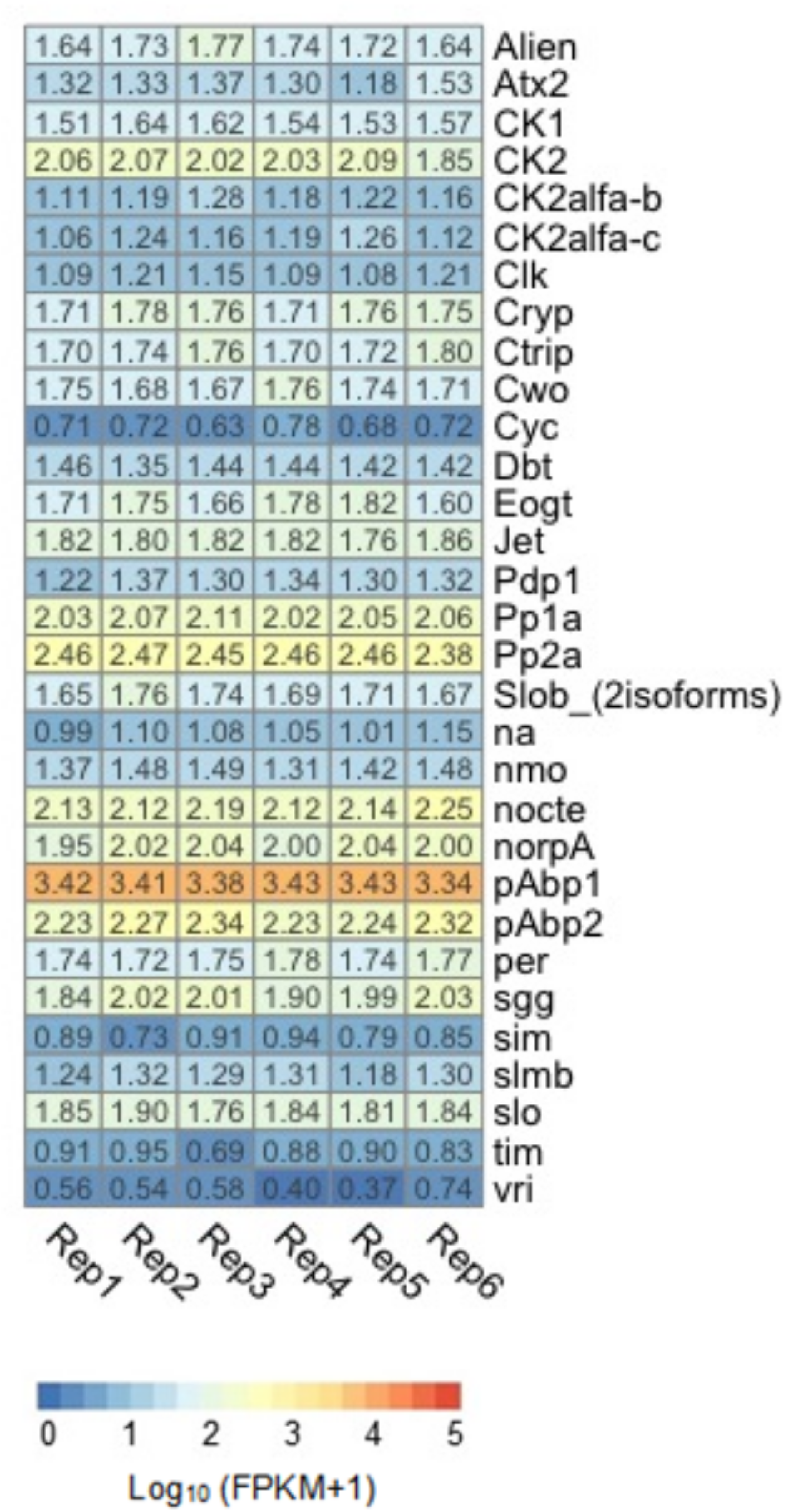
Expression profiles of clock-related genes. Heat map depicting the expression level of clock-related genes in the brain of *R. prolixus* nymphs. Expression level (displayed as Log10 (FPKM+1)) is represented by means of a color scale in which blue/orange represent lowest/highest expression. Each column represents the expression of one library.

Other genes related to the control of insect behavior include *foraging* (*for*) whose expression was detected in our brain transcriptome (S1 Table). This gene encodes a cGMP-dependent protein kinase and plays an essential role in modulating food search in different species of insects, such as *D. melanogaster* (60–62), locusts (63), ants (64), honeybees (65), and social wasps (66). In *R. prolixus, for* has been shown to participate in the modulation of locomotory activity (4,67), and its expression in the brain and fat body changes depending on the nutritional status of the insect, increasing with starvation (67). Our transcriptomic data seem to reinforce its relevance in the brain of *R. prolixus*.

### Sensory-related genes

Chemosensory proteins (CSPs) and odorant binding proteins (OBPs) can bind, solubilize and transport hydrophobic molecules (68). The role of these transporters has been mainly studied in insect antennae and other sensory tissues where they bind odor molecules (68,69). Nevertheless, the expression of certain CSPs and OBPs has been reported for insect guts, testes, Malpighian tubules and salivary glands (68,69). For this reason, we decided to data mine our database to characterize whether representatives of these protein families showed expression in bug brains (S2 Fig). Interestingly, high expression levels of several OBP and CSP transcripts, such as those of *RproCsp3, RproCsp5, RproCsp7, RproObp1, RproObp3, RproObp11, RproObp20*, and *RproObp26*, were detected in the brain of *R. prolixus*.

*Drosophila melanogaster takeout* 1 (*to*1) gene (*DmelTo1*) has been related to the regulation of feeding behavior and locomotor activity, and its expression has been detected in various fly structures and tissues, including the head, fat body, crop, and antennae (70,71). *DmelTo1* also affects male courtship behavior (72). The role of these proteins has been poorly studied in insects other than Dipterans, even though to date the scarce evidence also points to behavioral roles in a locust and a moth (73,74). Regarding triatomines, the expression of *to* genes has been reported in the digestive tract (25) and antennae of *R. prolixus* (22) and *T. brasiliensis* (75). In our brain transcriptome, *RproTo1, RproTo2, RproTo4*, and *RproTo6* were highly expressed (S3 Fig), suggesting that they may have relevant behavioral functions at the central level, as observed for *D. melanogaster*.

### Final remarks

Expression datasets obtained using transcriptomes represent powerful tools to uncover molecular targets for functional studies. Furthermore, these data allow improving automatically predicted gene models of interest through the manual curation of sequences (21,22). Therefore, they also increase the chances of performing successful functional experiments based on more trustable gene models. A drawback associated with whole tissue transcriptomes is that complex structures like the brain can present an intricate organization with specialized areas having very differentiated functional roles, and consequently, specific gene expression profiles.

Therefore, brain transcriptome studies should acknowledge that expression profiles represent averages of neuropiles having differentiated properties. This is especially true for clock genes or NPGs which can be expressed in very restricted sets of neurons. Therefore, any lack of differential expression observed in studies comparing levels of expression in different developmental or physiological conditions should be later validated with tissue-specific or single-cell sequencing methods, when available. Still, the high levels of expression on which we decided to focus here seem to denote relevant functions that deserve attention, as they might guide research toward specific targets allowing the development of more rational control methods.

---

**S1 Fig.**
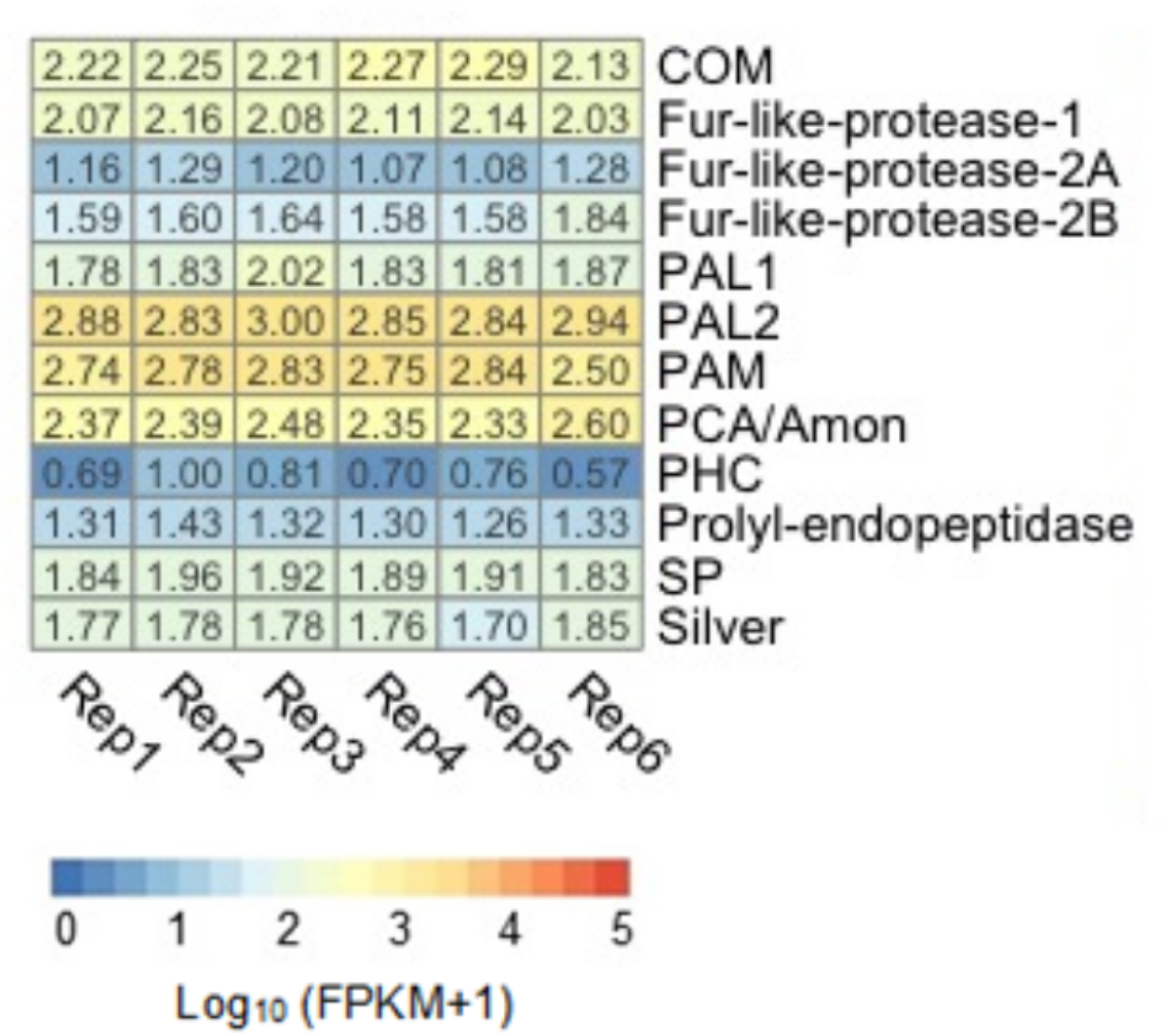
Expression profiles of neuropeptide processing enzymes genes. Heat map depicting the expression level of neuropeptide processing enzymes genes in the brain of *R. prolixus* nymphs. Expression level (displayed as Log10 (FPKM+1)) is represented by means of a color scale in which blue/orange represent lowest/highest expression. Each column represents the expression of one library.

**S2 Fig.**
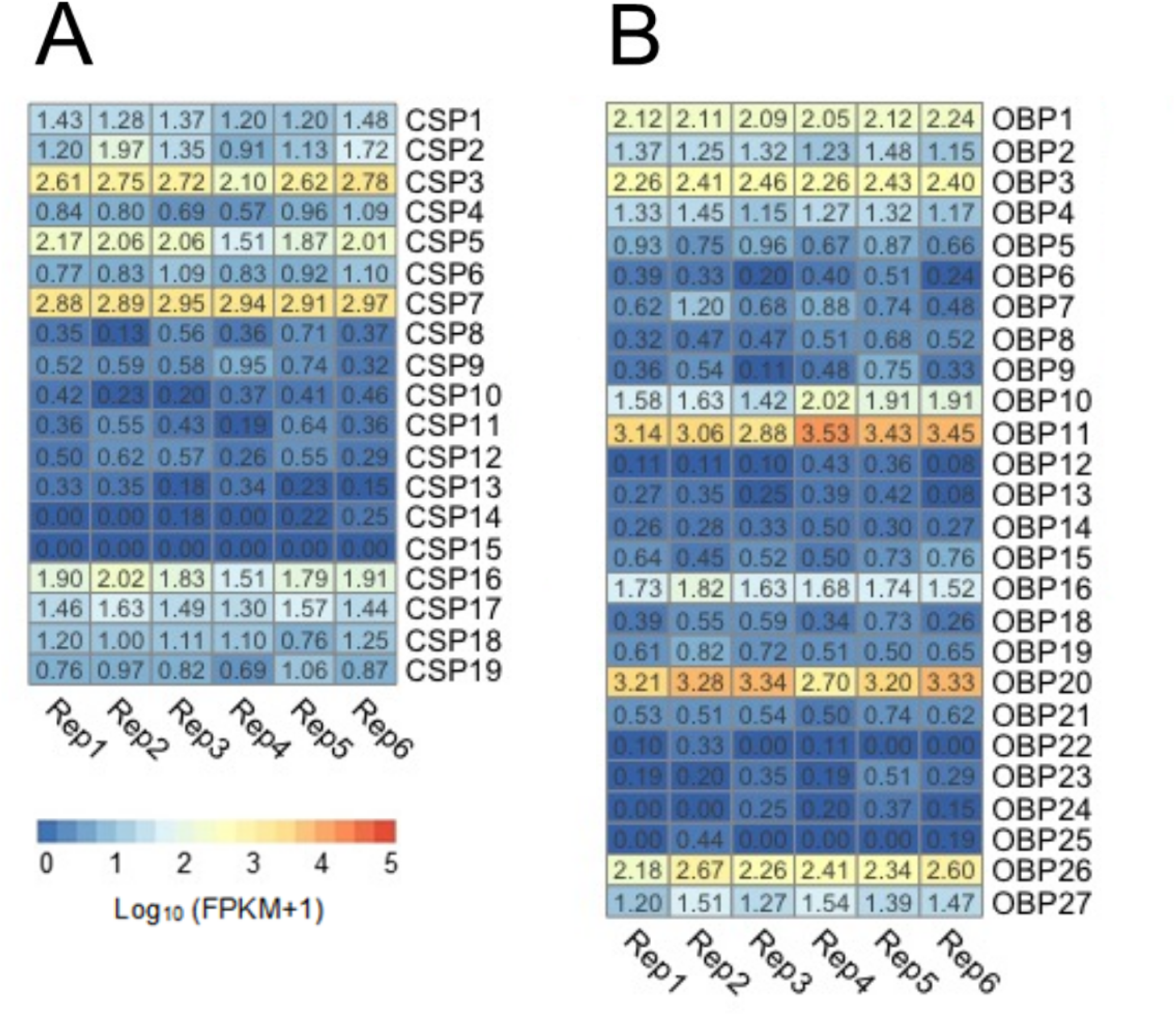
Expression profiles of chemosensory proteins and odorant binding proteins. (A) Heat map depicting the expression level of chemosensory proteins (CSPs), and (B) odorant binding proteins genes in the brain of *R. prolixus* nymphs. Expression level (displayed as Log10 (FPKM+1)) is represented by means of a color scale in which blue/orange represent lowest/highest expression. Each column represents the expression of one library.

**S3 Fig.**
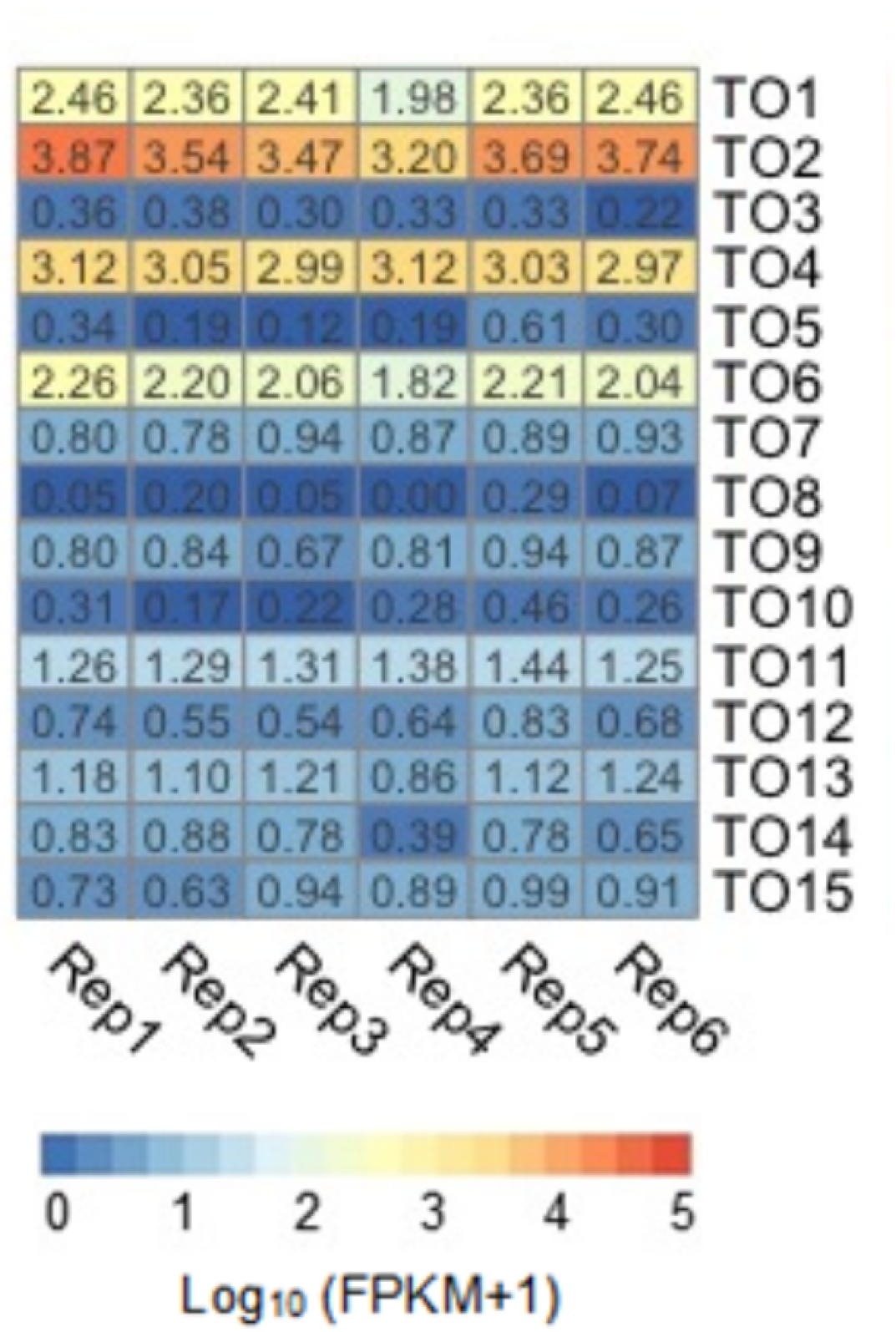
Expression profiles of takeout genes. Heat map depicting the expression level of takeout genes in the brain of *R. prolixus* nymphs. Expression level (displayed as Log10 (FPKM+1)) is represented by means of a color scale in which blue/orange represent lowest/highest expression. Each column represents the expression of one library.

**S1 Table. Details of the mRNA expression of Figs 1 – 4, S1 Fig, and S2 Fig**.

Columns are: the abbreviation of the gene assigned; gene name according to the annotation; VectorBase code–the official gene number in the RproC3 genome assembly; values of FPKM in each library (replicate); values of (Log10 (FPKM+1)) in each library. ND: not determined.

